# Profiling the effect of nafcillin on HA-MRSA D592 using bacteriological and physiological media

**DOI:** 10.1101/2020.04.30.070904

**Authors:** Yara Seif, Saugat Poudel, Hannah Tsunemoto, Richard Szubin, Michael J. Meehan, Connor A. Olson, Akanksha Rajput, Geovanni Alarcon, Anne Lamsa, Nicholas Dillon, Alison Vrbanac, Joseph Sugie, Samira Dahesh, Jonathan M. Monk, Pieter C. Dorrestein, Rob Knight, Joe Pogliano, Adam M. Feist, Bernhard O. Palsson, Victor Nizet

**Author notes:** **Correspondence:** To whom correspondence should be addressed: Victor Nizet, University of California, San Diego, 9500 Gilman Drive, La Jolla, CA 92093.

## Abstract

*Staphylococcus aureus* is a leading human pathogen associated with both hospital-acquired and community-acquired infections. The bacterium has steadily gained resistance to *β*-lactams and other important first-line antibiotics culminating in its categorization as an urgent threat by the U.S. Centers for Disease Control and Prevention. Observations of a varying response to antimicrobial exposure as a function of media type has revealed that clinical susceptibility testing performed in standard bacteriological media might not adequately represent pharmacological responses in the patient. Such observations have encouraged research designed to identify media types that more closely mimic the *in vivo* environment. In this study, we examine the response of a hospital-acquired USA100 lineage methicillin-resistant, vancomycin-intermediate *S. aureus* (MRSA/VISA) strain (D592) to nafcillin in a bacteriological compared to a more physiological tissue culture-based medium. We performed multi-dimensional analysis including growth and bacterial cytological profiling, RNA sequencing, and exo-metabolomics measurements (both HPLC and LC/MS) to shed light on the media-dependent activity of the commonly prescribed *β*-lactam antibiotic nafcillin.

|  |  |
| --- | --- |
| Design type(s) | • replicate design • transcription profiling design • sequence analysis objective |
| Measurement(s) | • minimal inhibitory concentration • cDNA • transcription profiling assay • time-course exo metabolome • cellular morphology • bacterial growth • Genome sequence |
| Technology Type(s) | • RNA sequencing • fluorescence microscopy • liquid chromatography-tandem mass spectrometry • high performance liquid chromatography • optical density measurement • Nanopore MinION • Illumina Hiseq 4000 |
| Factor Type(s) | • culture medium • biological replicate • experimental condition |
| Sample Characteristic(s) | • *Staphylococcus aureus* • culturing environment |

## Background and Summary

*Staphylococcus aureus* is a major pathogen that colonizes approximately 30% of the human population ^1^. Clinical *S. aureus* infections span a broad range of syndromes including skin and soft tissue infections, bacteremia, pneumonia, endocarditis, sepsis and toxic shock syndrome. Current antibiotics used to treat *S. aureus* target its cell wall, ribosome, nucleic acids and cell division machinery ^2^. The development of resistance to a large range of antibiotics including methicillin and other *β*-lactams, erythromycin, daptomycin, ciprofloxacin and others has placed *S. aureus* in the serious threat pathogens category. Vancomycin is a commonly prescribed intravenous medication for serious infections with resistant *S. aureus* strains, but an alarming emergence of vancomycin-intermediate and -resistant *S. aureus* strains (VISA and VRSA) highlights an urgent need for alternative treatment strategies.

Vancomycin binds to the D-Alanyl-D-Alanine residue of lipid murein precursors (UDP-MurNAc-pentapeptides) to disrupt downstream peptidoglycan assembly ^3^. *S. aureus* vancomycin resistance can be mediated by the *vanA* operon through hydrolysis of the D-Alanyl-D-Alanine precursors (vanX) and synthesis of D-Alanyl-D-Lactate dipeptides (vanAH), to which vancomycin does not bind ^3^. While our knowledge of the molecular basis for intermediate vancomycin resistance remains incomplete, mutations in *walKR* and *graRS* two-component regulatory systems are implicated in this phenotype ^4,5^. Decreased vancomycin susceptibility is also attributed to the high-inoculum effect, in which vancomycin binds to false targets, namely the D-alanyl-D-alanine residue in the peptidoglycan layer ^6^. As such, vancomycin is depleted in the peptidoglycan layer before it reaches its true target in the cytoplasm, an effect that is exacerbated as cell numbers increase. An alternative hypothesis postulates that the extensive use of vancomycin to treat methicillin-resistant strains predisposes them to step-wise polygenic mutations in genes encoding molecules involved in cell wall biosynthesis, increasing the probability of vancomycin resistance ^3^.

Here, we study a hospital-acquired USA100 lineage VISA strain, D592, isolated from a patient prior to vancomycin and daptomycin treatment ^7,8^. We examine the sensitivity of D592 to nafcillin in two different media types: 1) cation-adjusted Mueller-Hinton Broth (CA-MHB), the standard growth medium for antimicrobial testing in the clinical laboratory whose composition is poorly defined ^9^, and 2) Roswell Park Memorial Institute 1640 (RPMI), a chemically-defined standard mammalian cell culture medium better mimicking physiological conditions *in vivo ^10,11^*. However, *S. aureus* strains do not reliably grow on RPMI alone, despite the fact that RPMI contains all nutrients required to support growth *in silico* ^12,13^. Therefore, MIC was tested in RPMI supplemented with 10% Luria Bertani broth (RPMI+10%LB). Minimal growth requirements for *S. aureus* were recently uncovered ^13^. However, transcriptional regulation plays an important role in coordinating metabolic activity^14^ and results in condition-specific metabolic requirements ^15^. Therefore, the inability of *S. aureus* strains to grow on RPMI alone could be a result of transcriptional regulation.

We previously reported a similar data set for strain D712, which was collected from the same patient after daptomycin treatment ^16^. *S. aureus* strains exhibited different nafcillin susceptibility in these two media types ^10^. This data set was thus generated to study media-specific antibiotic susceptibility and how it is influenced by various environmental factors. Here, we investigate the response to nafcillin of VISA strain D592 in both bacteriological (CA-MHB) and physiological media (RPMI+10%LB). Between these two media the MIC value decreased from 256 **μ**g/ml in CA-MHB to 1 **μ**;g/ml in RPMI+10%LB. We measured the effect of nafcillin at various sub-inhibitory concentrations using optical density measurements, bacterial cytological profiling (BCP), transcriptomic sequencing (RNAseq), and high throughput exo-metabolomics (HPLC and LC/MS) experiments. BCP was performed to identify cytological parameters that change in response to antibiotic treatment, while RNAseq and metabolomics were used to measure changes in gene expression and secreted metabolites in response to the presence of nafcillin.

## Methods

The methodology used to generate this data set re-adapted from and similar to that used for our previously published articles ^16,17^.

### Culture and Growth Conditions

Standard bacteriological media MHB (Sigma-Aldrich) was supplemented with 25 mg/L Ca^2+^ and 12.5 mg/L Mg2+ (CA-MHB). Eukaryotic cell culture media RPMI 1640 (Thermo Fisher Scientific) was supplemented with 10% LB (R10LB). Broth microdilution was performed to determine the nafcillin MIC in each media condition. On the day of the experiment, overnight cultures of HA-MRSA D592 were diluted to a starting OD600 of 0.01 into fresh media and grown at 37°C with stirring to OD600 0.4. This preculture was then diluted back to OD600 0.01 into fresh media containing no drug or sub-inhibitory concentrations of nafcillin relative for each media type. Growth was monitored by obtaining OD600 readings every 45 min for 6 h. Three biological replicates were collected for the study, each derived from different colonies and overnight culture. Growth curves for all culture conditions are shown in **Fig. 1**.

**Figure 1.**
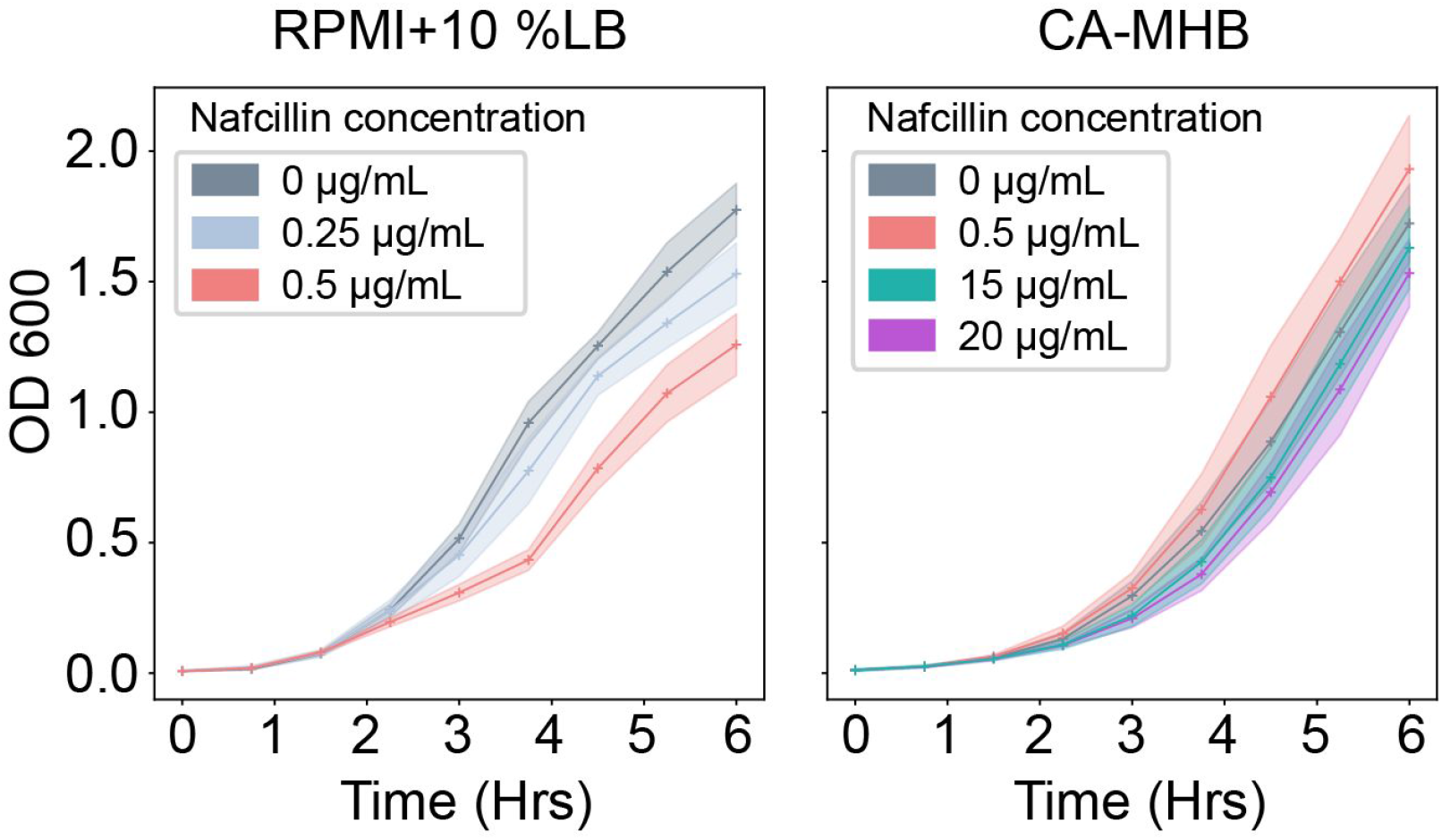
Growth curve for *Staphylococcus aureus* D592 strain in presence of nafcillin at various sub-inhibitory concentrations in CA-MHB and R10LB media.

### Bacterial Cytological Profiling

At the 3 h mark, samples were taken for fluorescence microscopy as previously described with minor modifications ^18–20^. In brief, 8 μL cells were added to 2 μL dye mix containing 10 μg/mL DAPI, 2.5 μM SYTOX Green, and 60 μg/mL FM4-64 in 1x T-base. The sample was then transferred to a glass slide containing an agarose pad (20% media, 1.2% agarose) and imaged on an Applied Precision DV Elite epifluorescence microscope with a CMOS camera. The exposure times for each wavelength were as follows, TRITC/Cy-5 = 0.025s, FITC/FITC = 0.01s, DAPI/DAPI = 0.015s, and were kept constant for all images.

Deconvolved images were adjusted using FIJI (ImageJ 1.51w) and Adobe Photoshop (2015.1) to remove background in WGA and DAPI channels and to ensure that cell and DNA objects are within the highest intensity quartile. These images were then processed using a custom CellProfiler 3.0 pipeline that individually threshold and filtered WGA and DAPI channels to obtain segmentation masks for the cell wall, DNA and entire cell. Objects identified in this manner were further processed in CellProfiler to obtain a total of 5,285 features ^21,22^. Prior to analysis, feature selection was necessary to create a subset of relevant features so as to minimize redundancy within the dataset. The summary of processing steps is presented in **Fig. 2**.

**Figure 2.**
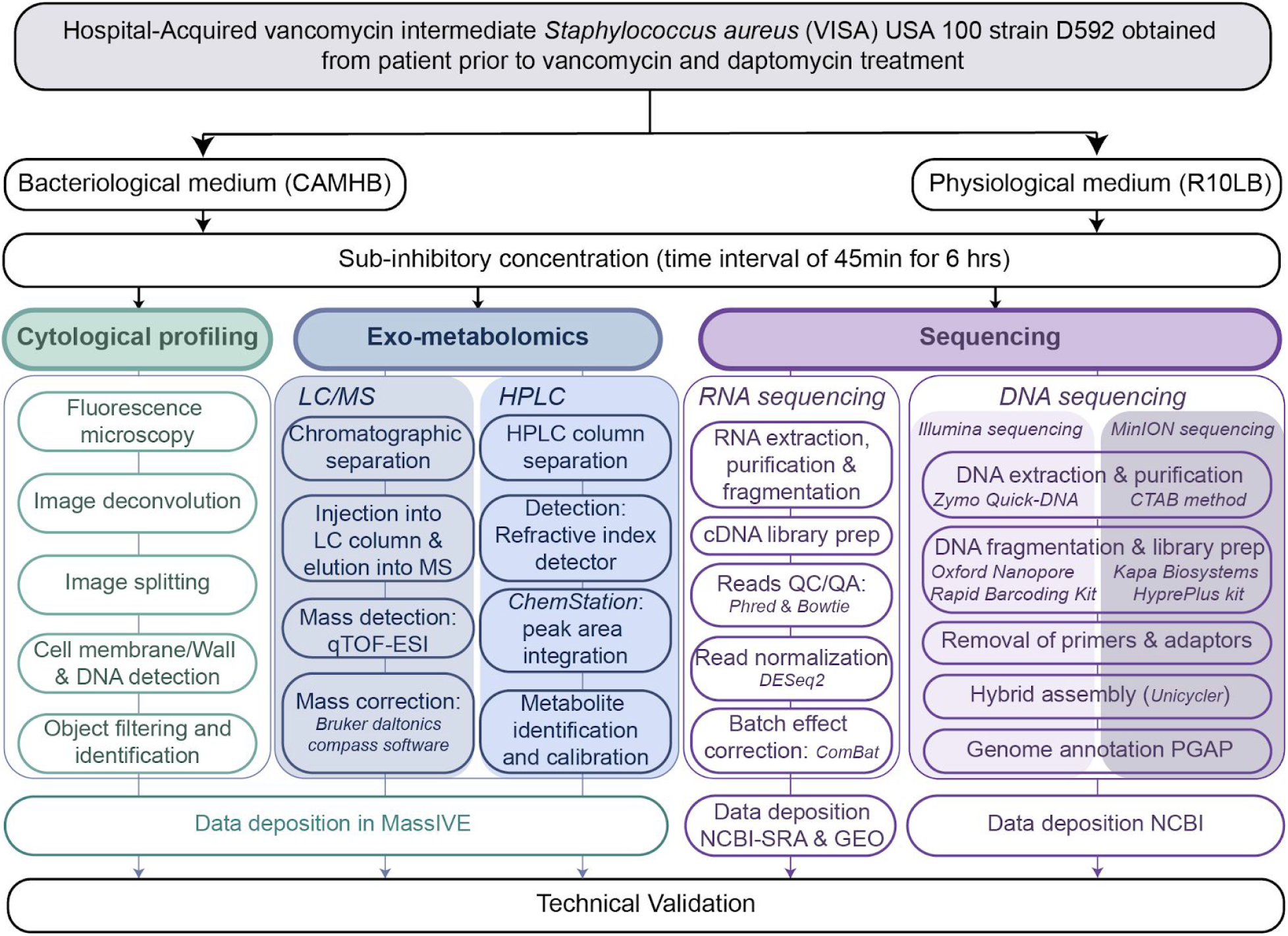
Diagram depicting the methodology of high throughput approaches used to profile the response of *Staphylococcus aureus* strain D592 to nafcillin in bacteriological and physiological media.

### DNA sequencing and genome assembly

The reference genome of D592 was sequenced using an Illumina Hiseq 4000 (paired end, 100/100 bp reads) and Nanopore MinION to 50x coverage. For Illumina sequencing genomic DNA was prepared using a Zymo Research Quick-DNA Fungal/Bacterial Microprep Kit, and libraries were prepared using a Kapa Biosystems HyprePlus kit. For MinION sequencing, high molecular weight genomic DNA was prepared using a CTAB method and libraries were prepared using an Oxford Nanopore Rapid Barcoding Kit. Prior to assembly, the quality control steps were performed to remove unincorporated primers, adaptors, and detectable PCR primers. The genome was then assembled into 2 contigs (genome and plasmid) using Unicycler 0.4.2 in “default” mode for hybrid assemblies and annotated using the NCBI Prokaryotic Genome Annotation Pipeline (PGAP) v4.11. The sequencing reads show an average Phred quality score of >32, which corresponds to a base calling accuracy of 99.99%. The reads were submitted to NCBI under accession number SRP258107 and the complete record was deposited at NCBI as NZ_CP035791.1. The final chromosomal genome size was 2,820,117 base pairs, while the final plasmid size was 27,267 base pairs.

### cDNA library preparation and RNA sequencing

After 3 hours of growth (at mid-log phase), 3 mL samples were taken for RNA sequencing and added to tubes containing 6 mL RNAprotect. After incubation, they were centrifuged to remove the supernatant. RNA was extracted from the pelleted cells using a ‘Quick RNA Fungal/Bacterial Microprep’ kit developed by Zymo Research. Cells were mechanically lysed with a Roche MagNa Lyser instrument and DNA was removed with DNase I during the RNA purification. RNA quality was checked with an Agilent Bioanalyzer instrument and ribosomal RNA was removed using an Illumina Ribo-Zero kit. Remaining RNA was used to build a cDNA library for sequencing using a KAPA Stranded RNA-seq Library Preparation Kit. The kit was used for RNA fragmentation, sequencing adapter ligation, and library amplification. Generated cDNA libraries were sent for Illumina sequencing on a HiSeq 4000.

### RNA sequencing analysis

The phred quality scores for illumina sequencing were generated using Fastqc package^23^. Bowtie2 was used to align the raw reads to the D592 genome and to calculate alignment percentage, FastQC^24^. The aligned reads were then normalized to transcripts per million (TPM) with DESeq2. The ComBat module within the sva package of R was used to correct for batch effects ^25,26^. Distance matrices for hierarchical clustering were calculated with sklearn package^27^. The summary steps are provided in **Fig. 2**.

### Untargeted Liquid Chromatography Mass Spectrometry Data Acquisition

Following dilution of the preculture of HA-MRSA USA100 D592 into fresh media, approximately 400 μL of liquid media containing cells were collected at 45 min intervals (at the same time as samples for OD600 measurements) from each of the samples. Growth media was syringe-filtered through 0.22 μm disc filters (Millex-GV, MilliporeSigma) to remove cells. The filtered growth media was collected and stored at −80 °C until analysis by liquid chromatography mass spectrometry (LC/MS). For LC/MS analysis, samples were subjected to chromatographic separation using an UltiMate 3000 UHPLC system (Thermo Scientific). Chromatographic separations were achieved using a 50 mm × 2.1 mm Kinetex 2.6 micron polar-C18 column (Phenomenex) held at a fixed temperature of 30 °C within an actively heated column compartment. Samples were injected onto the LC column via thermostatted autosampler maintained at 4 °C. For samples containing RPMI + 10% LB media the injection volume was 5 μl, while the injection volume was 2 μl for samples containing CA-MHB to prevent excessive oversaturation of the mass spectrometer detector due to the higher concentrations of many molecules in the CA-MHB media.

After injection, the sample components were eluted from the LC column into the mass spectrometer using a flow rate of 0.5 mL/min and the following mobile phases: Mobile phase A was LC/MS grade water with 0.1% formic acid (v/v) and mobile phase B was LC/MS grade acetonitrile with 0.1% formic acid (v/v). The LC gradient program was as follows: 0–1.0 min 5%B, 1.0–5.0 min 5–35%B, 5.0–5.5 min 35–100%B, 5.5–6.0 min 100%B, and 6.0–6.5 min 100–5%B followed by 5 minutes of re-equilibration at 5%B. Mass spectrometric data was acquired using a Bruker Daltonics maXis Impact quadrupole-time-of-flight (qTOF) mass spectrometer equipped with an Apollo II electrospray ionization (ESI) source and controlled via otofControl v4.0.15 and Hystar v3.2 software packages (Bruker Daltonics). The mass accuracy of the maXis instrument was first externally calibrated using a calibration solution of sodium formate which provided >21 reference m/z’s between 50–1500 m/z of the mass spectrum in both positive and negative polarities (reference m/z list provided within instrument control software).

Sodium formate solution was prepared using 9.9 ml of 50/50% isopropanol/water, plus 0.2% formic acid, and 100 μl of 1 M NaOH. During infusion of all samples, the mass accuracy of the instrument was maintained to <10 ppm *via* constant introduction of an internal calibrant, or “lock mass”, in the form of hexakis (1H,1H,2H-difluoroethoxy)-phosphazene (SynQuest Labs, Inc.). During positive polarity runs the lock mass compound was detected as the ion m/z 622.028960 (C12H19F12N3O6P3+) and in negative polarity the lock mass compound formed the ion m/z 556.001951 (C10H15F10N3O6P3−).

Instrument source parameters were set as follows: nebulizer gas (Nitrogen) pressure, 2 Bar; Capillary voltage, 3,500 V; ion source temperature, 200 °C; dry gas flow, 9 L/min. The global mass spectral acquisition rate was set at 3 Hz. The instrument transfer optics were tuned as follows: Ion funnel 1 & 2 RFs of 250 Vpp (volts peak-to-peak), hexapole RF of 100 Vpp, quadrupole ion energy of 5 eV, collision quadrupole energy of 5 eV, and a TOF pre-pulse storage of 7.0 μsecs. The post-collision quadrupole RF and TOF transfer time were stepped across four values per MS scan. The collision RF was stepped at 450, 550, 800, 1100 Vpp. The transfer time was stepped at 70, 75, 90 and 95 μsecs. All samples were run twice, once under positive polarity settings and once under negative polarity settings.

Following acquisition of the LC/MS data, lock mass calibration was applied to all data files to apply a linear correction calibration to all m/z values recorded in each mass spectrum. The application of this mass correction was applied automatically via the Bruker Daltonics Compass Data Analysis software (ver. 4.3.110), using the m/z of the hexakis (1H,1H,2H-difluoroethoxy) phosphazene as the reference lock mass calibration compound. Following lock mass re-calibration of the data, all files were converted from the Bruker Daltonics proprietary format (.d) and exported to an open data format known as .mzXML. All data herein was deposited to MassIVE ^28^. The brief methodology is provided in **Fig. 2**.

### Targeted High-Performance Liquid Chromatography

For organic acid and carbohydrate detection, samples were collected every 45 min and filtered as described above. The filtered samples were loaded onto a 1260 Infinity series (Agilent Technologies) high-performance liquid chromatography (HPLC) system with an Aminex HPX-87H column (Bio-Rad Laboratories) and a refractive index detector. The system was operated using ChemStation software. The HPLC was run with a single mobile phase composed of HPLC grade water buffered with 5 mM sulfuric acid (H2SO4). The flow rate was held at 0.5 mL/min, the sample injection volume was 10 μL, and the column temperature was maintained at 45 °C. The identities of compounds were determined by retention time comparison to standard curves of acetate, ethanol, glucose, lactate, pyruvate, and succinate. The peak area integration and resulting chromatograms were generated within ChemStation and compared to that of the standard curves in order to determine the concentration of each compound in the samples. These final concentration values were deposited to MassIVE database^28^. The procedure of HPLC is depicted in **Fig. 2** and measured concentrations for three carbon sources are shown in **Fig. 3**.

**Figure 3.**
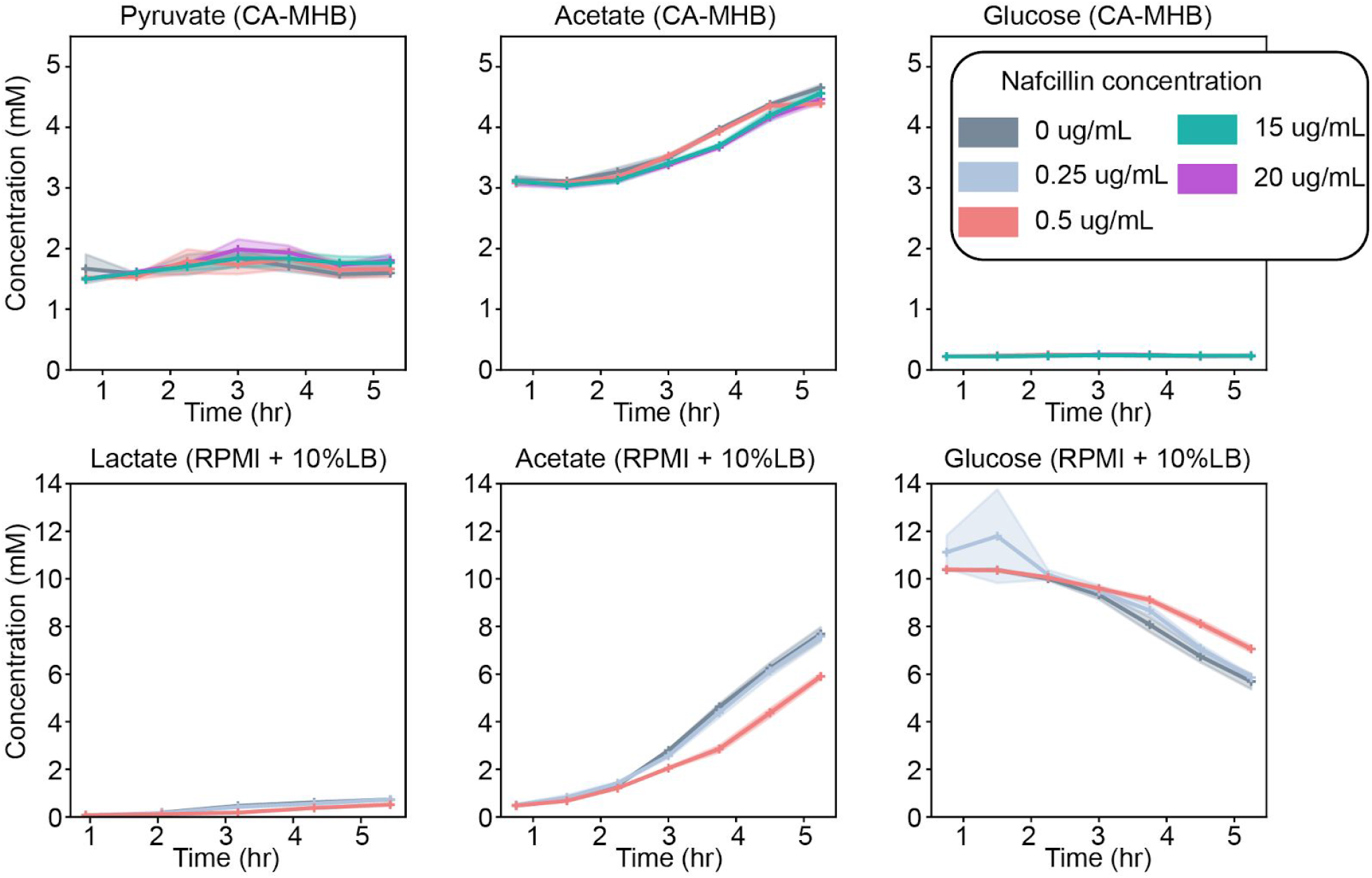
HPLC-derived quantitative time-course exo-metabolomics measurements for S*taphylococcus aureus* D592 cells exposed to various antibiotic concentrations in RPMI + 10%LB and CA-MHB. Here, we show the absolute calibrated concentrations of acetate and D-glucose in both media types, pyruvate in CA-MHB and lactate in RPMI + 10%LB.

## Data Records

The reference genome assembly for *S. aureus* D592 was submitted in NCBI under accession number NZ_CP035791.1 and the corresponding reads were submitted under accession number SRP258107. The optical density and the corresponding calculated growth rates as well as the minimal inhibitory concentration measurements are available on Figshare ^29,30^. The BCP, HPLC, and Mass spectrometry data have been deposited to MassIVE repository under accession number MSV000085316 ^28^. The complete RNAseq pipeline can be found on Figshare ^31^, Fastq files for each run have been deposited on NCBI-Sequence Read Archive under accession number PRJNA526576 ^32^. The overall summarized statistics of RNAseq are available on Figshare ^33^. The metadata for all multi-omics data types are available on Figshare ^34^.

## Technical Validation

### Bacterial Cytological profiling

The technical validation of BCP was done through manual screening during the image segmentation process in CellProfiler. Accurate cell and object traces and measurements were verified manually for representative images. The cell outlines were matched with corresponding related structures e.g., DNA through “parent” tags. Finally, the output files for the cellular features were uploaded to MassIVE repository^28^. A representation of the image analysis pipeline for BCP data is provided in **Fig. 4**.

**Figure 4.**
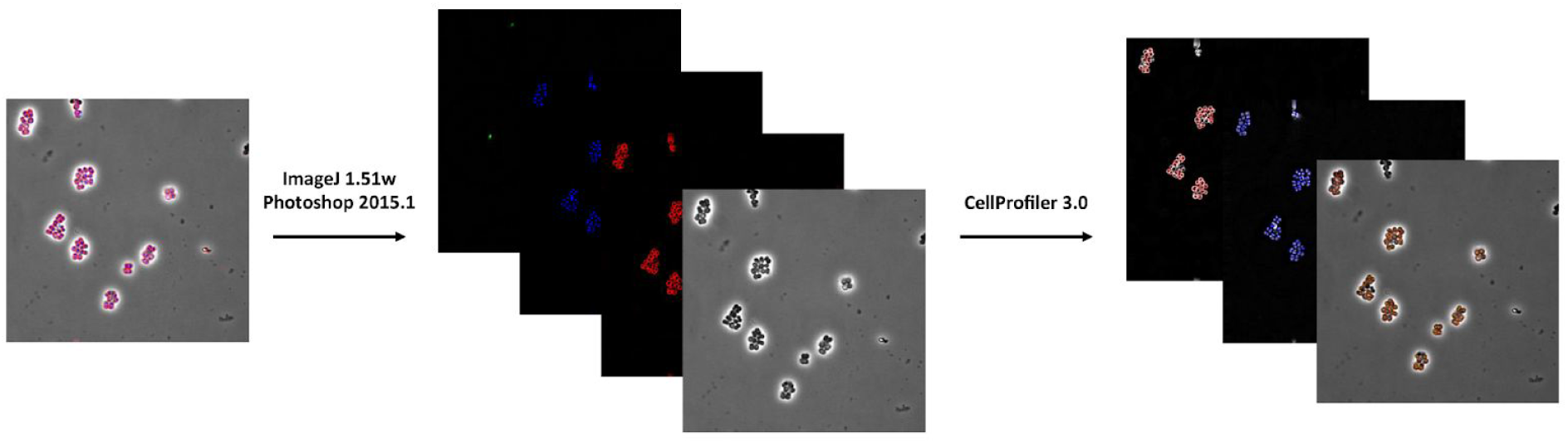
Depiction of image analysis pipeline for Bacterial Cytological Profiling (BCP) of *Staphylococcus aureus* strain D592.

### Untargeted Liquid Chromatography Mass Spectrometry data acquisition

For each sample, the base peak chromatogram (BPCs) and multiple extracted ion chromatograms (EICs) were compared to evaluate the reproducibility of global retention time and ion intensity. The reproducibility of retention time and peak intensity were obtained by comparing the BPCs of each experimental triplicate. The EICs of the molecules were evaluated using retention time drift and peak area of <0.1 min and <15% correspondingly.

### RNA sequencing

The alignment of reads with the reference genome gives an average alignment score of 97.47%. Because the number of samples was large, their processing was done in three different batches. Batch effects were corrected using the ComBat module of the SVA package in R. As a result, expression profiles showed a Spearman’s correlation coefficient >0.975 between biological replicates. The RNAseq results are shown in **Fig. 5**.

**Figure 5.**
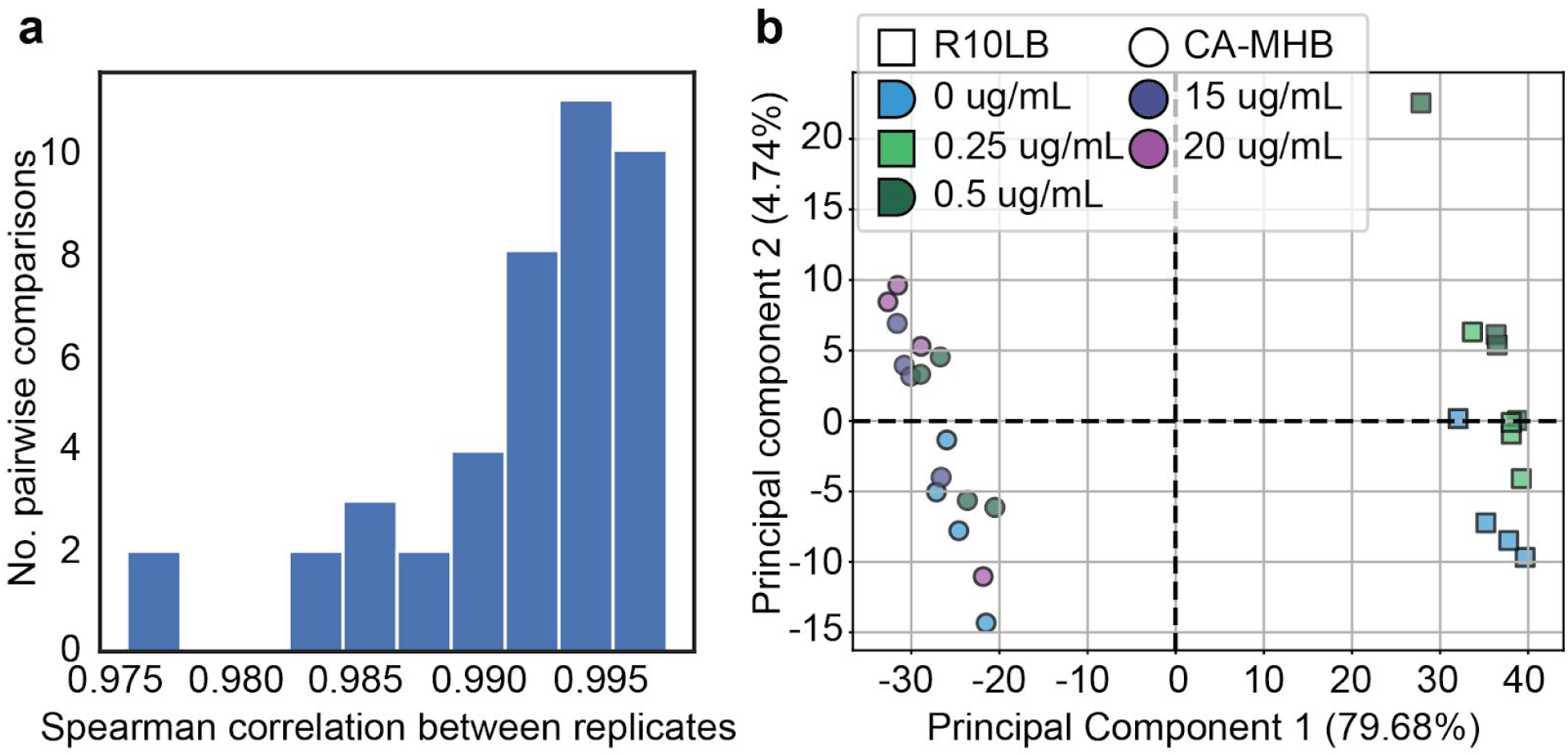
Quality control for transcriptomics sequencing data. **a)** Histogram of Spearman's correlation coefficient for TPM values between biological replicates. **b)** PCA plot for expression profiles across all samples. Samples on the left of the plot represent cultures grown in CA-MHB, while samples to the right of the plot represent cultures grown on RPMI +10%LB. Each sample point is color coded by nafcillin concentration to which the cultures were exposed and shaped according to the culture media type.

## Code availability

The complete RNAseq pipeline used in analysis of RNAseq data is available on Figshare ^31^. The script to remove batch effects from RNAseq data is also available on Figshare ^35^.

## Acknowledgements

We thank Anand Sastry for helping build the RNA sequencing analysis pipeline and Marc Abrams for reviewing this manuscript. This research was supported by NIH NIAID grant (1-U01-AI124316).

## Author Contributions

Y.S. compiled, analyzed results and wrote the Data Descriptor, Methods and Technical Validation, S.P. analyzed the RNA sequencing data and wrote Methods, H.T. performed growth experiments, analyzed the BCP data, and wrote Methods subsections, A.R. reviewed the article, M.M. analyzed the HPLC data and wrote subsections of the Methods, R.S. prepared samples for RNA sequencing and libraries for DNA sequencing and wrote Methods subsections, C.A.O. prepared samples for HPLC and wrote Methods subsections, A.L. performed growth experiments and analyzed BCP data, N.D. performed preliminary growth and MIC experiments. A.V. performed growth experiments. J.S. wrote Methods. S.M.D. performed growth experiments. B.O.P and V.N. conceptualized the project, obtained funding, and edited the final manuscript.

